# Network resonance during slow-wave sleep facilitates memory consolidation through phase-coding

**DOI:** 10.1101/565242

**Authors:** Quinton M. Skilling, Brittany C. Clawson, Bolaji Eniwaye, James Shaver, Nicolette Ognjanovski, Sara J. Aton, Michal Zochowski

**Affiliations:** Biophysics Program, University of Michigan, 930 N University Ave., Ann Arbor, MI 48109; Department of Molecular, Cellular, and Developmental Biology, University of Michigan, 830 N University Ave., Ann Arbor, MI 48109; Applied Physics Program, University of Michigan, 450 Church St, Ann Arbor, MI 48109; Department of Physics, University of Michigan, 450 Church St, Ann Arbor, MI 48109

**Keywords:** Memory Consolidation, Phase coding, Oscillatory dynamics, Sleep, Resonance

## Abstract

Sleep plays a critical role in memory consolidation, however, the exact role that sleep and its effects on neural network dynamics play in this process is still unclear. Here, we combine computational and experimental approaches to study the dynamical, network-wide underpinnings of hippocampal memory consolidation during sleep. We provide data to support a novel hypothesis on the role of cellular resonance with sleep-associated theta band (4-12 Hz) hippocampal oscillations in this process. We show that increases in the stability of hippocampal memory representations after learning (which predicts successful memory consolidation) are mediated through emergent network-wide resonance and locking of neuronal activity to network oscillations. These changes arise in the network as a function of changes to network structure during learning, and mirror experimental findings in the hippocampus. Finally, we show that input-dependent pattern formation (e.g. “replay”) in the hippocampus during sleep states, together with spike timing dependent plasticity (STDP)-based memory consolidation, leads to universal network activity reorganization. This reorganization generates heterogeneous changes in neuronal spiking frequency, similar to what has been observed in a variety of brain circuits across periods of sleep. Our results support the hypothesis that sleep plays an active role in memory consolidation by switching the hippocampal network from rate-based to phase-based information representation. The mechanisms through which this occurs supports the integration of heterogeneous cell populations into memory traces.

**Significance Statement:** In this study, we provide a mechanistic explanation of how sleep selectively facilitates memory consolidation, through recruitment of heterogeneous neuronal populations and structural reorganization of the network into an engram. Specifically, we show that emergent theta band oscillations during sleep facilitate stabilization of memory representations via spike timing dependent reinforcement. This stabilization, together with STDP, allows for systematic reorganization of synaptic connections within these populations, universally redistributing firing rates of participating neurons. Simultaneously, network oscillations facilitate a switch from rate-to phase-coding of information among neuronal populations with highly heterogenous firing frequencies, incorporating more neurons into the engram. Our results reconcile discrepant findings on network reorganization during sleep, and demonstrate a clear mechanism for both strengthening and weakening of synaptic efficacy during sleep.

## Introduction

It is widely hypothesized that new information is encoded in brain circuits through activity-dependent, long-term synaptic structural changes (1), which serve as the physical scaffold for memory traces (or engrams) (2, 3). While certain features of information storage may be localized to specific cell populations (e.g., location information encoded in hippocampal place cell activity), in general, localizing memory traces to specific neural circuits has been an elusive task (2). Furthermore, recent data indicate that engrams are formed from neuronal populations which are vastly heterogeneous in terms of their firing patterns, with log-normal distributions of firing rates ranging over many orders of magnitude (4, 5). These populations appear to encode different aspects of experience (5–7) and may exhibit different dynamics during memory consolidation processes. A critical unanswered question is how these heterogeneous populations, distributed throughout the brain over vast synaptic distances, cooperate in the process of long-term memory storage.

A major hurdle for understanding the systems-level mechanisms for memory consolidation is that very little is known about the dynamical substrates required for long-term information storage in neural circuits and, conversely, what impact learning has on subsequent network dynamics. A widely-accepted hypothesis, supported by experimental evidence, is that new information is encoded in a network based on immediate, experience-associated changes in the activity of specific neurons (often referred to as “engram neurons”) (8). Such representation of information constitutes a mnemonic “rate code”. However, rate coding has limitations for long-term information storage in the brain *in vivo*. First, individual neurons have widely divergent baseline firing rates, as described above. This wide range of firing rates would constitute background “noise”, obscuring firing rate changes in a sparse subset of neurons representing new information. Second, because individual neurons have a fairly limited dynamic range over which their firing rates can vary, rate coding has a limited capacity for neuronal information integration over time. In other words, neurons’ peak firing rates would represent a “ceiling”, after which no additional information could be carried by a given neuron. Third, changes in firing rate will alter spike timing-dependent plasticity (STDP) rules governing activity-dependent synaptic strengthening or weakening between constituent network neurons. As neurons increase their firing rates, they will either reduce STDP-mediated synaptic strengthening/weakening (due to differences in firing rates between pre-and postsynaptic neurons), or bias STDP toward potentiation (9). In either case, the information-carrying capacity of STDP for the network is limited by firing rate increases. Thus, for long-term information storage (consolidation) in a network of heterogeneous neurons, non-rate-based coding mechanisms must be invoked.

Another long-accepted assumption is that following information encoding during learning, memories are consolidated, via either modification of existing synaptic connections or *de novo* creation of additional synapses. Such synaptic changes constitute a structural network heterogeneity, which subsequently serves as a so-called dynamical attractor in the network (10, 11). When such an attractor is present, incomplete patterns of spiking among engram-encoding neurons could lead to subsequent recollection of the complete pattern, i.e. a memory. This recollection process possibly consists of augmentation of a memory trace with new populations of cells that were not as active initially (5). It remains unclear what activity-dependent mechanisms could mediate consolidation leading to such an attractor formation. However, oscillatory patterning, such as that occurring during sleep, has been implicated in promoting synaptic plasticity (4, 5, 12–14). Various network oscillations appear preferentially in specific brain circuits (e.g., hippocampal and thalamocortical circuits) during particular behavioral states (i.e. waking, REM and NREM sleep) and have been established to be highly predictive of state-dependent cognitive processes and network plasticity (15). Further, network oscillations have been implicated in promoting STDP by precisely timing the firing between pairs of neurons (8, 14, 16). Thus, hypothetically, oscillatory dynamics could mediate a switch between rate coding (present during learning) and timing-dependent coding (in which neurons’ respective timing, rather than firing rate, carries information). We have recently shown that in a weakly coupled regime, an activity-dependent shift in theta-band cellular resonance among neurons may play an important role in explaining learning-initiated network reorganization, via STDP-based mechanisms (8). We hypothesize that, for the reasons outlined above, this switch from rate to phase coding (i.e. neurons’ respective phase of firing on each cycle conveys the information about the network patterning) is essential for long-term information storage in brain networks.

Here, we combine computational modeling of a highly reduced CA1 network with the analysis of *in vivo* mouse CA1 activity to elucidate the role of theta band (4-12 Hz) oscillations in short-and long-term reorganization of hippocampal network structure and dynamics during contextual fear memory (CFM) consolidation. Contextual fear conditioning (CFC) leads to rapid formation and consolidation of new memories (i.e. after single-trial learning) (17). CFM consolidation relies on *ad lib* sleep in the hours immediately following CFC (18, 19), and is associated with augmented theta-frequency activity in CA1 in the hours following CFC (20). Critically, recent *in vivo* work has shown CFM is disrupted when theta oscillations are suppressed during post-CFC sleep via optogenetic or pharmacogenetic inhibition of CA1 fast-spiking interneurons (20). Conversely, CFM consolidation can be rescued from disruption caused by experimental sleep deprivation when theta oscillations are driven optogenetically (via rhythmic activation of fast-spiking interneurons) in CA1 (18).

We show that resonance properties of the CA1 network model are sufficient to recreate **all of these** experimental phenomena, including the state-dependence of fear memory consolidation, disruption of consolidation through interneuron inhibition, and rescue of consolidation when network oscillations are artificially driven. We demonstrate that STDP-based memory consolidation during a dynamical network state analogous to NREM sleep: (1) causes increased stability of functional network connectivity patterns, (2) requires augmentation of theta band oscillations like those observed during post-learning sleep, and (3) causes dramatic, differential changes in the activity profile of highly active vs. sparsely firing neuronal populations. These same dynamic changes are not observed during the dynamical network state corresponding to waking. Together, these results show that successful memory consolidation requires the brain to switch between information coding schemes which occur naturally through the sleep-wake cycle. In so doing, the functional network structures associated with engrams become more stable and leads to successful long-term memory storage.

Finally, similar frequency dependent changes in firing frequency mediated by NREM sleep, were observed in other modalities than the hippocampus (i.e. visual cortex [5]). Thus we conclude that even though this study is limited to investigating role of theta frequency network oscillations in fear memory, it may yield insight into potential universal neuronal network mechanisms of information storage in the brain.

## Results

### Introduction of memory traces augments network oscillatory dynamics and stability of functional connectivity patterns

To investigate the mechanisms involved in sleep-dependent memory consolidation, we simulated a reduced CA1 network model composed of two cell types: excitatory pyramidal neurons and inhibitory interneurons that loosely represent parvalbumin positive (PV+) interneurons. For cells in the model, we used a conductance-based formalism (see **Methods**) incorporating a slow-varying potassium current which acts as a control parameter for neural spiking dynamics (21). In the brain this conductance is regulated by muscarinic acetylcholine (ACh) receptors (22). During wake, high ACh blocks these pyramidal cells’ slow-varying potassium currents, yielding increased gain in the firing frequency response to varying (excitatory) input and decreased capacity for firing to synchronize with rhythmic input (collectively referred to as type 1 excitability). Low levels of ACh, seen during NREM sleep, allows these slow-varying potassium currents to play a larger role in membrane excitability. In this scenario, the model’s pyramidal neurons exhibit spike frequency adaptation, reduced spike frequency response gain, and increased synchronization capacity – features characteristic of what is referred to as type 2 excitability (21). Hence, in this model, sleep/wake dynamics are controlled by switching pyramidal neurons from type 2 membrane excitability (sleep) to type 1 excitability (wake). In contrast, PV+ interneurons in the model consistently exhibit type 1 dynamics.

The switch from type 1 to type 2 pyramidal cell excitability results in a modification of neurons’ responses to excitatory input – from being integrators, where firing frequency is the main carrier of information, to resonators with a slowly increasing input-frequency curve, where the spike frequency distributions are significantly narrower. A reduction in firing frequency distribution at the transition from wake to NREM sleep is observed in numerous brain structures including the hippocampus and cortex *in vivo (23)*. In this resonating regime, neurons have an increased capacity to synchronize to periodic input, and phase-of-firing relationships (rather than firing frequency relationships) play prominent roles in carrying information. Here, when the network is weakly coupled, we find that an oscillatory pattern emerges spontaneously through resonance of the pyramidal and interneuron populations, consistent with prior findings (14). Within this framework, we first investigated how changes in excitability in subsets of neurons, an analogy for memory encoding by “engram neurons”, affect network activity patterns. To this end, we mimicked the formation of an engram by artificially increasing the outgoing synaptic connections of a randomly chosen subset of 250 pyramidal neurons by various degrees, which we refer to as the “memory strength”. Comparing raster plots for the network in the NREM sleep state (type 2 dynamics), before vs. after introduction of the engram (Figure 1A-B; corresponding to a memory strength of 20), reveals the emergence of well-defined oscillations and, correspondingly, an increase in theta-band spectral power (4-12 Hz; Figure 1C), consistent with previous work (18). The emergence of network oscillations with phase locking of neuronal spiking (i.e. characteristic oscillations in the theta-band regime) might be sufficient to generate reliable sequences of spiking patterns; such oscillation-driven sequences are thought to play an important role in memory consolidation. To investigate this fully, we calculated the relative stability of network functional connectivity, using the functional network stability metric (FuNS; see **Methods**) (24).

**Figure 1:**
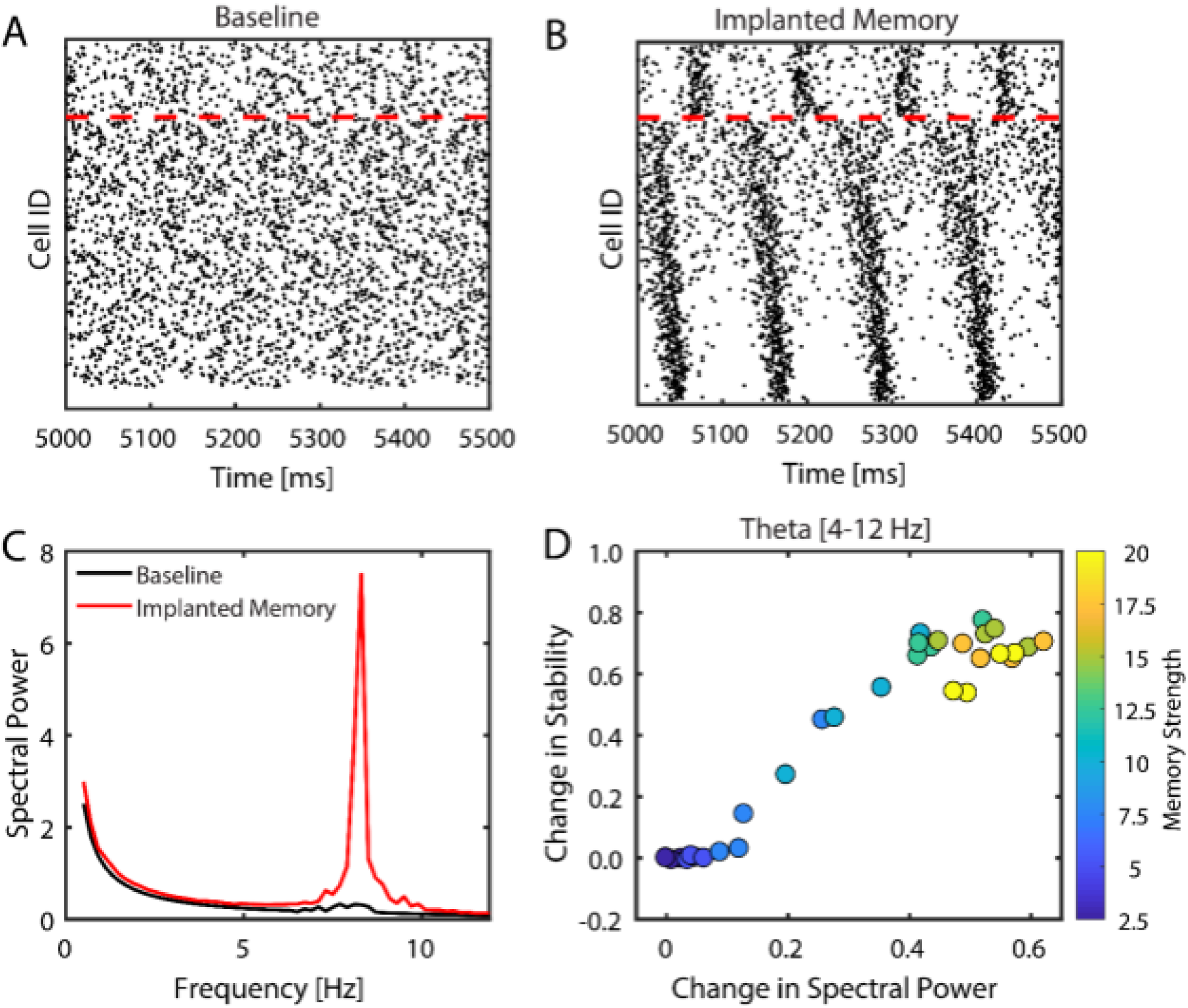
Model networks respond to sparse strengthening of excitatiory synapses through emergance of theta rhythm and phase-locking. **A-B)** Raster plots of the network before (left) and after (right) introduction of a memory of strength 20. Neurons below the red, dashed line are excitatory and those above the line are inhibitory. Inhibitory neurons connected randomly to 50% of the other inhibitory neurons and to 30% of the pyramidal neurons whereas excitatory neurons connected randomly to 6% of both the excitatory and inhibitory neurons; a random subset of excitatory neurons increased their synaptic strength to other excitatory neuron, constituting the memory strength. **C)** Fourier transform of the excitatory network signal reveals a sharp increase in spectral power only near 8 Hz. **D)** Measuring the change in stability as a function of the change in spectral power density (integrated over theta-band frequencies) for increasing the strength of those selected synaptic connections reveals a linear releationship. The color bar represents the magnitude of synaptic strength change from baseline.

The FuNS metric is sensitive to reconfiguration of functional connectivity patterns within the network, but not to fluctuations in the precise spike order between pairs of neurons. High stability can be detected for weak but consistent functional connectivity patterns. Therefore, FuNS qualitatively provides information about temporal evolution of functional connectivity as compared to direct assessment of correlation between spike bouts.

By increasing the memory strength, we observed a positive linear relationship between increased FuNS and increased theta-band power in the network (Figure 1D). This suggests that locking of neuronal firing with network oscillations, and associated stabilization of network functional connectivity patterns, is driven by synaptic potentiation between engram neurons and other neurons in the network. This agrees with previous experimental results, showing that contextual fear memory consolidation is associated with both increased theta-band oscillations, and increased FuNS, during sleep in the hours immediately following conditioning (17, 18, 20). We proceed to show that it is this phase-locking of neurons to underlying oscillations that is important for memory consolidation.

### Hippocampal network stabilization in vivo predicts effective memory consolidation

We hypothesized that network plasticity in hippocampal area CA1 following single-trial contextual fear conditioning (CFC) (25) would be a plausible biological system to investigate how rapid memory encoding affects subsequent neural network dynamics. Since CA1 network activity is necessary for fear memory consolidation in the hours following CFC (26), we recorded the same population of CA1 neurons over a 24-h baseline and for 24 h following CFC to determine how functional network dynamics were affected by *de novo* memory formation. C57BL/6J mice implanted with bundles of CA1 stereotrodes either underwent CFC (placement into a novel environmental context, followed 2.5 min later by a 2-s, 0.75 mA foot shock; n = 5 mice), sham conditioning (placement in a novel context without foot shock; Sham; n = 3 mice), or CFC followed by 6 h of sleep deprivation (a manipulation known to disrupt fear memory consolidation [18, 19, 27]; SD; n = 5 mice) (Figure 2). Spike data from individual neurons was discriminated offline using standard methods (consistent waveform shape and amplitude on the two stereotrode wires, relative cluster position of spike waveforms in principle component space, ISI ≥ 1 ms) (5, 17, 18, 20, 28). Only neurons that were stably recorded and reliably discriminated throughout the entire baseline and post-conditioning period were included in subsequent analyses of network dynamics.

**Figure 2:**
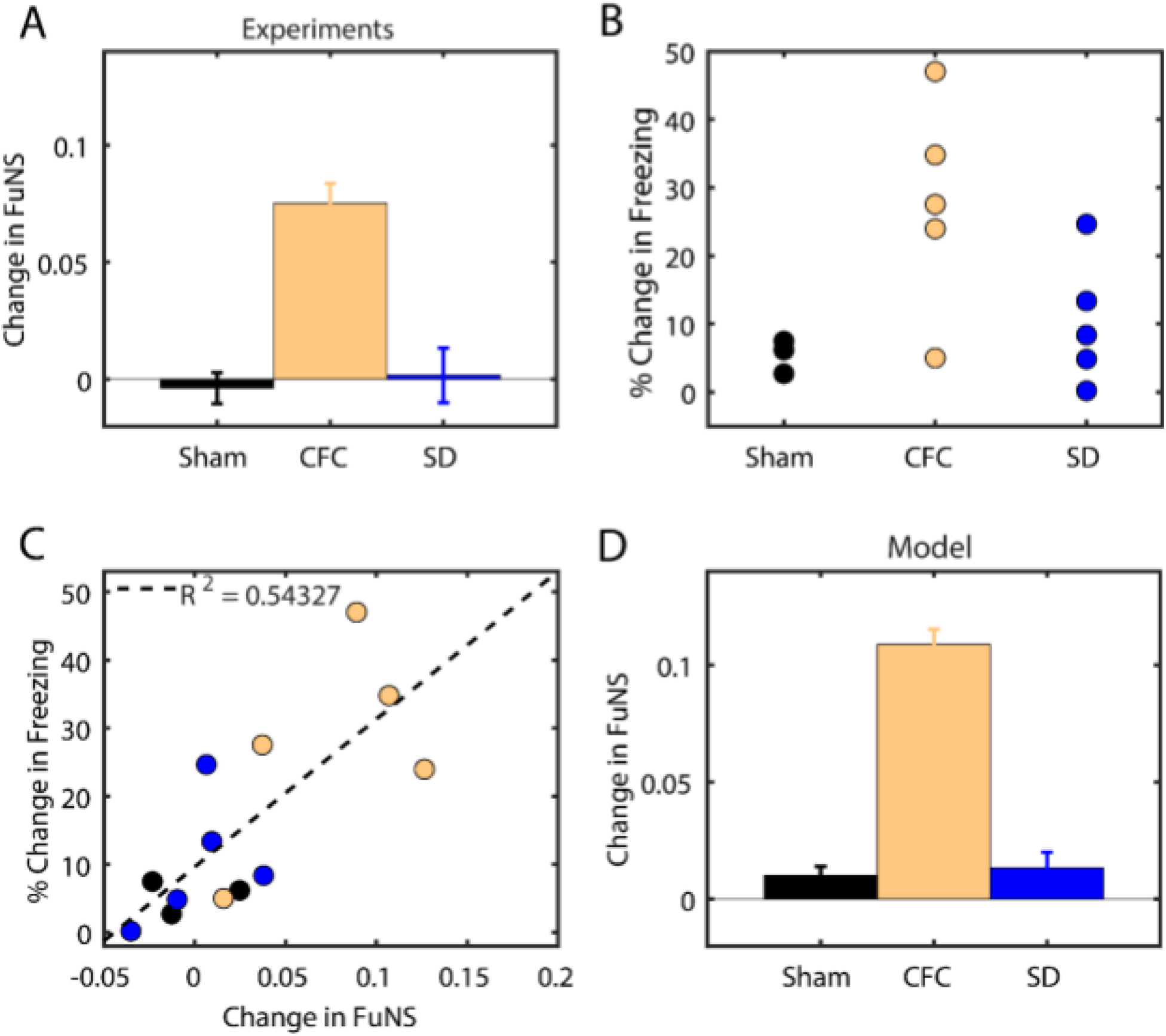
Functional network stability predicts future level of memory consolidation. **A)** Analysis of *in vivo* recordings in the mouse hippocampus CA1 area following CFC for corresponding behavioral states (sham — black [n = 3], SD — blue [n = 5], CFC – Gold [n = 5]) reveals that ad lib sleep post conditioning leads to the greatest increase in FuNS. **B)** Percent change in freezing, behavioral correlate of learning, taken 24 hours after exposure to a fear stimulus. **C)** Percent change in freezing as a function FuNS changes observed during the 24 hour priod. **D)** Model predictions for the change in FuNS in each simulation group: Sham (learning in SWS states without selective connectivity strenghtening; n= 5) and SD (learning in Wake dynamical state with selective connectivity strenghtening; n = 5) show only marginal changes in FuNS whereas CFC (learning in NREM dynamical state with selective connectivity strenghtening; n = 5) show a maximal increase in FuNS. All error bars represent the standard error of the mean.

We measured changes in FuNS in these recordings after each manipulation by quantifying FuNS on a minute-by-minute basis over the entire pre-and post-training (i.e., 24 h duration) intervals and calculating their respective difference within each animal. Consistent with previous findings (17), we observed a significant increase in FuNS over the 24 h following CFC during NREM sleep (Figure 2A). In contrast, no change in NREM FuNS was seen in Sham mice or following CFC (during recovery sleep) in SD mice. Group differences in NREM FuNS were reflected in the behavior of the mice 24 h post-training, when context-specific fear memory was assessed (Figure 2B). Mice allowed *ad lib* sleep following CFC showed significantly greater freezing behavior when returned to the conditioned context than in Sham and SD mice. Moreover, training-induced changes in NREM-specific FuNS for individual mice predicted context-specific freezing during memory assessment (Figure 2C). Thus, successful consolidation of a behaviorally-accessible memory trace *in vivo* is accompanied by increased FuNS in the CA1 network.

### Network stabilization is driven by memory encoding, oscillatory dynamics, and STDP

To better assess the network requirements for memory consolidation, we returned to our model network and allowed synapses between pyramidal neurons to undergo spike timing-dependent plasticity (STDP, see **Methods**), while we monitored changes in FuNS. We simulated networks with differential exposure to learning (e.g., CFC) and varying vigilance states to mimic the experiments described above. Briefly, the Sham condition was modeled as cycling through Wake-NREM-Wake (type 1 – type 2 – type1) without an encoded memory (i.e., a memory strength of 1). The CFC condition was modeled as cycling through Wake-NREM-Wake with the network having a memory strength of 10. Finally, the SD condition was also modeled as a network with a memory strength of 10, but subsequently, neuronal excitability parameters associated with wakefulness (i.e. type 1 excitability) were maintained. We found that only in the CFC condition was the network capable of successfully consolidating information, maintaining stable network dynamics over time (Figure 2D). In both Sham and SD conditions, the network representations showed no significant increase in stability, even in the context of STDP-based learning. Based on these results, we hypothesized that theta-frequency network oscillations, augmented by type 2 networks with memory strengths greater than one, are critical for successful STDP-based consolidation of a hippocampal memory trace. We focus on this phenomenon in the following sections.

### Theta-band resonance in the network mediates increased network stability

Recent *in vivo* work has shown the importance of parvalbumin-expressing (PV+) fast-spiking interneurons in hippocampal fear memory consolidation (18, 20, 29). Optogenetic or pharmacogenetic inhibition of PV+ interneurons in hippocampal area CA1 decreased theta-band oscillations during both post-CFC REM and NREM sleep, without altering the firing frequency of neighboring cells. These manipulations are sufficient to disrupt contextual fear memory consolidation (18, 20). Conversely, fear memory can be rescued from disruptive effects of experimental sleep deprivation when PV+ interneurons are rhythmically activated at theta frequency (18), at which the CA1 network naturally resonates (20). Based on prior computational work (14), we hypothesized that this frequency, and not others, is effective for memory rescue because of natural subthreshold oscillatory theta-band frequency exhibited by pyramidal neurons.

To examine this further, we manipulated the inhibitory neurons in our model networks with an established encoded memory (i.e. increasing heterogeneity connection strength 10 fold), by (a) silencing them (see **Methods**), and then (b) driving their firing to occur periodically, at variable frequencies (via driving them with 2ms DC current pulses at a given frequency) similar to what was done *in vivo* by Ognjanovski et al. (20). In Figure 3A, we show that silencing inhibitory neurons following introduction of an encoded memory eliminated the change in network stability. However, stimulation at 8 Hz was sufficient to restore FuNS. Further, the peak in theta-band spectral power typical of CFC modeling experiments is diminished when inhibitory neurons are silenced but returns under 8 Hz stimulation (Figure 3B). At the same time, similarly to the experiments, there was no overall increase in the spiking frequency of the network’s pyramidal neurons (Figure 3C), which would be expected from a PING-type (rather than resonance-based) mechanism. Importantly, varying the frequency of the applied stimulation to the inhibitory population showed that at frequencies near 8 Hz, across the network there were peak measures of (a) spike-field coherence (Figure 3D), (b) FuNS (Figure 3E), and (c) maximum spectral power (Figure 3F; see **Methods**), similarly to what was observed *in vivo* (20). This provides a clear indication that resonance characteristics of neuronal excitability, amplified in type 2 dynamics of NREM sleep, lead to phase-locking behavior across the network following memory encoding. Indeed, example raster plots in Figure 3G allow visual inspection that well-defined phase locking occurs only near the 8 Hz stimulation frequency.

**Figure 3:**
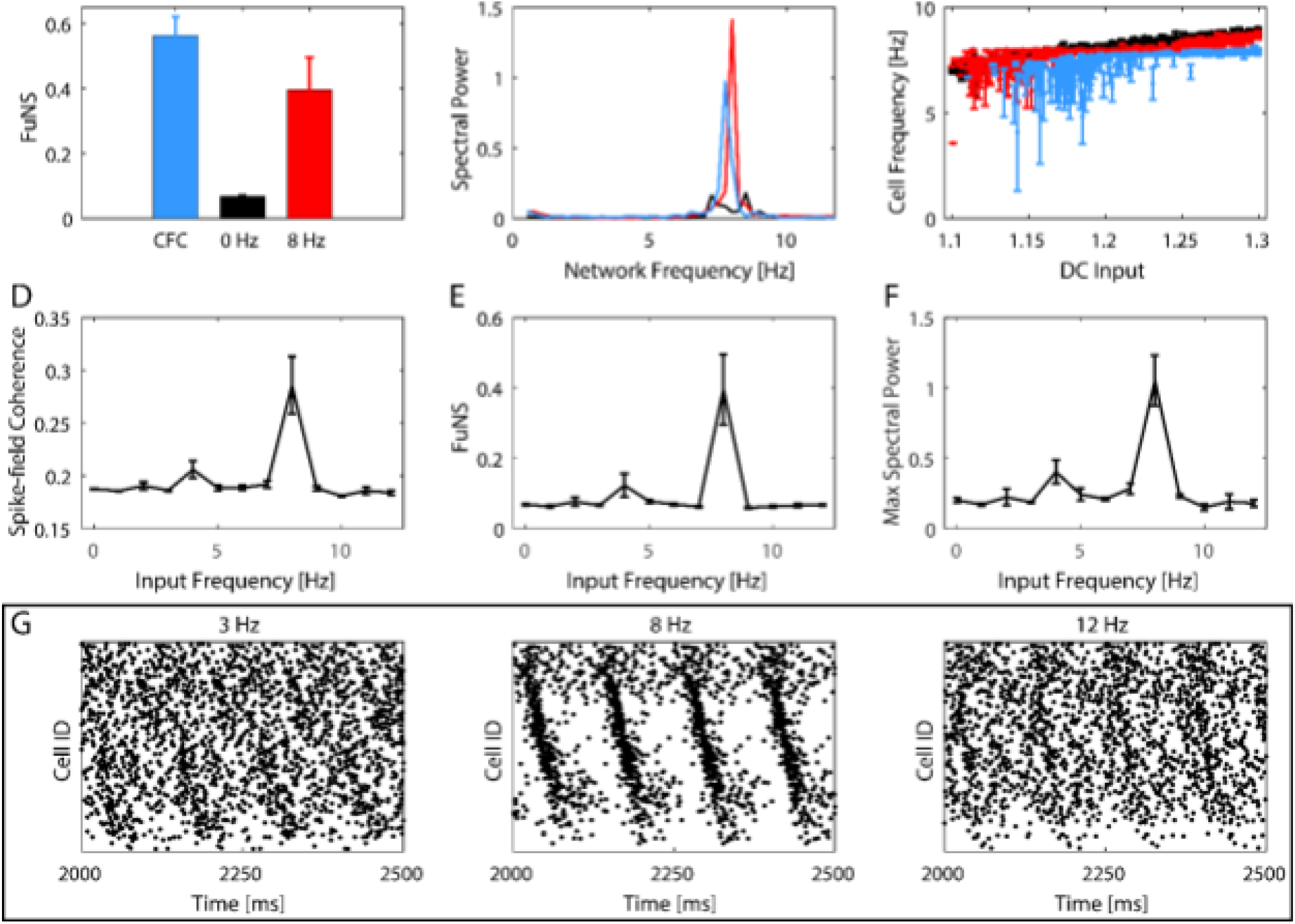
Optogenetic silencing/driving of inhibitory interneurons reveals network-level resonance. **A)** Comparisons of FuNS for standard CFC dynamics (blue), when inhibitory cells are silenced (black) and when inhibitory cells are driven with a 2 ms wide pulse at 8 Hz. **B)** Spectral power as a function of network frequency for the same cases as above. **C)** Individual neuron frequency as a function of input drive into that neuron. All colors are conserved across the top three panels. **D)** Spike-field coherence, **(E)** FuNS and **(F)** the maximum spectral power as a function of pulse frequency applied to silenced inhibitory neurons. **G)** Example raster plots of pyramidal neurons without strengthened connections for driving silenced inhibitory neurons at 3 Hz (left), the resonant 8 Hz (middle), and 12 Hz (right). All errorbars represent the standard error of the mean for n = 3.

### Input-dependent phase locking of firing to network oscillations predicts firing rate reorganization across a period of sleep

A series of recent studies have demonstrated that sleep has heterogeneous effects on neuronal firing rates within neural circuits. Specifically, initially highly active neurons reduce their firing rates across a period of sleep, while sparsely firing neurons simultaneously increase their firing rate (4, 5, 30). Sleep is essential for this redistribution of firing rates, as firing rate changes do not occur in animals that are experimentally sleep deprived (5).

We hypothesized that this phenomenon could result from input-dependent phase-locking of neurons’ firing to NREM sleep oscillations. Specifically, we predict that neurons that are highly active during wake would fire at an earlier phase than sparsely-firing neurons during subsequent NREM sleep oscillations. Figure 4 illustrates the relationship between neurons’ phase of firing (calculated with regard to the inhibitory network signal; see **Methods**) calculated during NREM sleep before learning as a function of the normalized frequency during pre-learning wake. We observed that the activation of fastest firing neurons during waking indeed occurs earlier in the phase of the pyramidal network oscillation during sleep. This suggests that neurons take on a phase coding strategy during network oscillations (i.e., those occurring during sleep) which reflects relationships in the prior rate coding strategy present during wake. Based on this relationship, we hypothesized that excitatory connections from high-firing to low-firing neurons are strengthened via STDP during sleep, while connections from low-firing to high-firing neurons are weakened.

**Figure 4:**
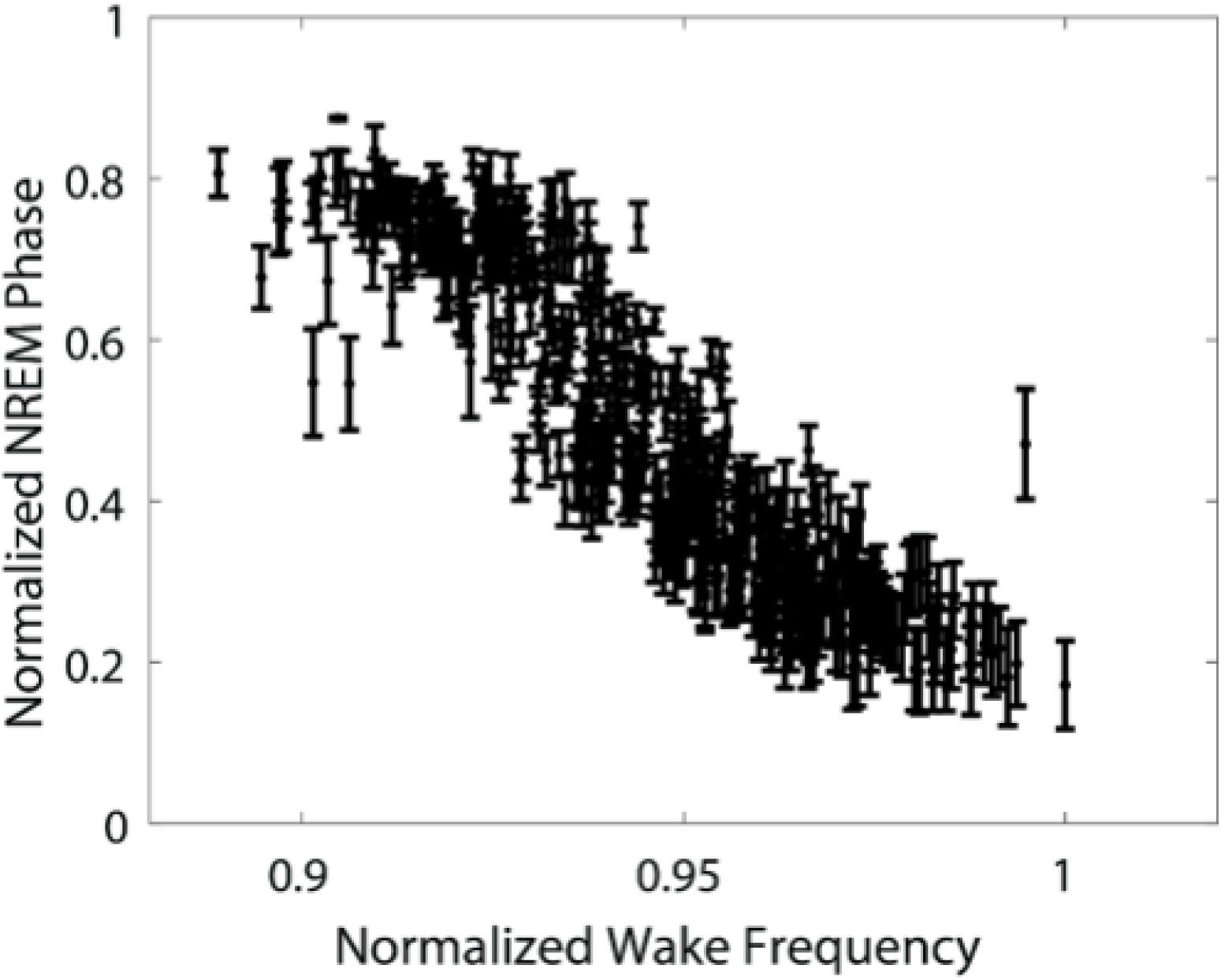
Phase of firing in NREM versus wake frequency of pyramidal neurons reveals that the neurons firing with the highest frequency align with an earlier phase of the pyramidal network population whereas slower firing neurons align with later phases.

To test this, we examined how individual neuron frequencies change during STDP-based memory consolidation. In our network model, we compared the effects of consolidation of an encoded memory during NREM sleep (similar to DMSO mice *in vivo*) with (a) a condition where network oscillations are abolished (by silencing inhibitory neurons, as in Figure 3 and similar to CNO mice *in vivo*) and (b) a condition where the network is in a continuous wake-like state (i.e., sleep deprivation, SD, where theta-band oscillations are less pronounced). We examined the changes in wake-state neuronal firing frequencies after a consolidation interval in each of these scenarios. In the model, when STDP-based memory consolidation took place during sleep (without manipulations to the inhibitory network, DMSO), we observed a significant increase in the firing rates of pyramidal neurons initially firing at low rates during wake; in contrast, those with the highest initial firing showed a decrease in firing rate (Figure 5A, **top**). By color-coding neurons based on their relative change in frequency across sleep, we identify in the corresponding pre-learning NREM raster plot (Figure 5A, **bottom**) that neurons that fire faster (slower) not only show a decrease (increase) in firing frequency due to learning, but also fire earlier (later) in the oscillation, consistent with Figure 4. Note that some neurons did not fire during NREM and so did not show a significant change in firing frequency due to learning (black neurons in Figure 5A, **top**). By comparison, silencing of inhibitory neurons in the network (CNO) led to a homogenous increase in firing rates for those neurons active consistently across learning (Figure 5B). Finally, when the network was maintained in a wake state (i.e. high Ach) following memory encoding (mimicking SD), there was no significant change in neuronal firing rates (Figure 5C). We compared these results with data recorded from the hippocampus of mice subjected to corresponding experimental manipulations (18, 20). We measured the change in log firing frequency across a six-hour time interval at the start of the animals’ rest phase (i.e., starting at lights on), with recordings occurring either the day before CFC (baseline) and immediately after training (post CFC). Following CFC, mice either were (a) allowed *ad lib* sleep with administration of a vehicle (DMSO), (b) were allowed *ad lib* sleep with pharmacogenetic inhibition of hippocampal PV+ fast-spiking interneurons (CNO), or (c) were sleep deprived. Changes in log firing rate for all stably-recorded neurons were calculated in each condition, as a function of their baseline firing rate. The resulting best-fit lines reveal that all cases show relatively low rescaling before exposure to fear but that initially low firing neurons increase their firing rate in response to fear conditioning, with the greatest increase seen in the best learners, DMSO mice (Figure 5D). Indeed, comparing the slopes of firing rate change (Figure 5E) for post-shock recordings show significantly weaker reorganization of firing rates in CNO and SD than in in DMSO condition. This rescaling of firing rates across the network is an important prediction of the model, as it suggests a possible universal network-level correlate of sleep-based memory consolidation *in vivo*.

**Figure 5:**
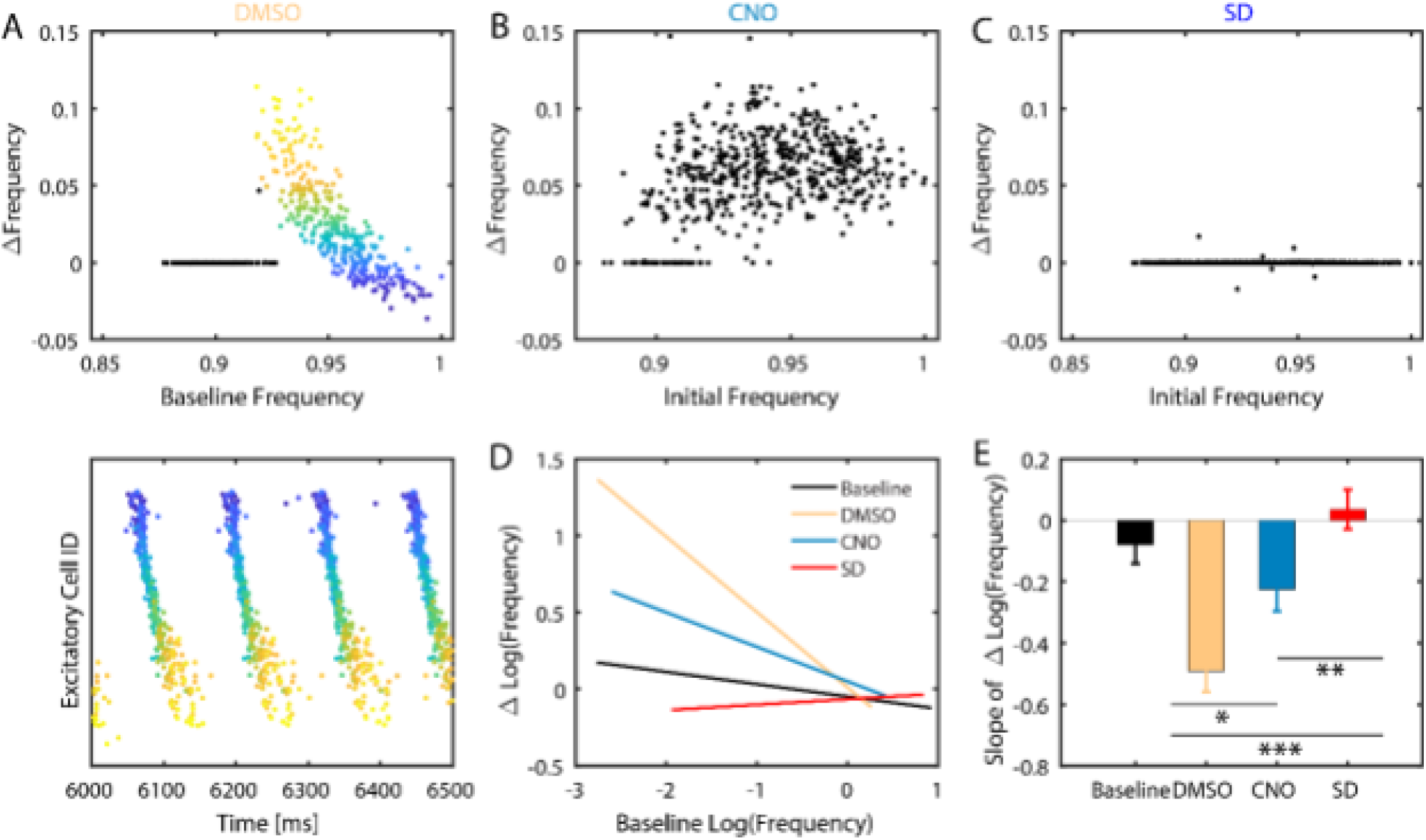
Memory consolidation during sleep differentially affects frequency of firing neurons – model prediction and experiment. MODEL: **A)** Changes in individual neuron firing frequency (normalized to baseline) due to memory consolidation during sleep as a function of normalized baseline firing frequency (top) and a snapshot of the corresponding raster plot in NREM sleep before learning. Neurons are color-coded based on their change in frequency from baseline and the color is conserved in the raster plot. Black-colored neurons (top) are those which did not fire during NREM sleep. Of the neurons that are consistently active, those with initially lower frequency increase their frequency whereas neurons with initially higher frequency decrease their frequency. **B)** Change in firing frequency (normalized to baseline) as a function of normalized baseline firing frequency. Unlike in DMSO, firing frequency changes homogenously across baseline firing rates. **C)** Conversely, no change in firing frequency (normalized to baseline) is observed in SD as a function of the normalized baseline frequencies. **EXPERIMENT**: **D)** Best fit lines of the change in log firing rates vs the initial log firing rate, comparing baseline recordings (solid lines; composite n = 11) to post recordings (dashed lines) for the first 6 hours post CFC in experimental DMSO (Yellow; n = 3), CNO (Teal; n = 3), and SD (Blue; n = 5). **E)** Slope comparison of change in log firing rates for DMSO, CNO, and SD baseline and post-shock recordings. Analysis of covariation revealed statistically significant slope differences between DMSO and CNO (*, p = 0.0114), CNO and SD (**, p = 0.0084), and DMSO and SD/Baseline (***, p < 0.0001).

## Discussion

Sleep has long been known to be vital for successful memory consolidation. Sleep’s requirement for long-term memory storage has been demonstrated across organisms and across different types of memories (e.g., those mediated by network activity in the hippocampus vs. sensory cortex)^15^. Similarly, recent advances have shown that varying oscillatory dynamics accounts for different aspects of memory consolidation (20, 31, 32). Here, we argue that neuronal phase-locking to network oscillations expressed during sleep promotes feed-forward synaptic plasticity (i.e., STDP) that promotes long-term memory storage.

Our models demonstrate that the initial encoding of memories in the hippocampus during CFC increases theta-band oscillation power. The augmentation of these oscillations occurs in a NREM sleep-like, low-ACh, type 2 network state, but not a high-Ach, wake-like type 1 state. Such oscillations mediate input-dependent phase locking of neuronal spiking, due to the neuronal excitability properties observed in the low ACh type 2 state. This locking to network oscillations leads to more stable firing relationships between neurons. We observe this increase in stability (as increases in FuNS), both in our hippocampal network model after introduction of a synaptically-encoded memory (Figure 1), and experimentally after hippocampus-dependent memory encoding *in vivo* (Figure 2). This change appears to result from stable mapping between frequency response during wake and relative phase pattern during NREM sleep (Figure 4). The network subsequently undergoes structural re-organization through classic STDP mechanisms, leading to renormalization of firing frequencies (Figure 5), with sparsely firing neurons increasing their firing rate substantially, and highly active neurons decreasing their firing rate. These results are consistent with experimental observations in CA1 during CFM consolidation (Figure 5), and observed firing rate changes in neurons across periods of sleep in other brain structures (4, 5).

By assessing the relative change in FuNS recorded in the CA1 network following CFC, we have determined the salient features of network wide dynamics that accompany successful sleep-dependent memory consolidation (Figures 2-5) (4, 5, 17, 18, 20, 23). We find that increased FuNS is not only a result of stronger oscillatory patterning of the network, but predicts whether experimental conditions will support, disrupt, or rescue fear memory consolidation (Figure 2).

Thus, taken together, our results hint at a universal mesoscopic network mechanism – emergence of spike-timing dependent FuNS – underlying what is commonly referred to as systems consolidation. The process of systems consolidation leads to formation of a widely-distributed engram from what is initially a transient, discrete, and localized group of synaptic changes (leading to increased firing in a sparse populations of neurons).

Thus, our results point to the hypothesis that while the brain may use a rate code to initiate memory encoding during experience, memories can only be consolidated through phase-based information coding, in the form of phase locking of firing network oscillations in subsequent sleep. This is supported by the idea that rate-based information coding in the brain is highly limited (as described in the introduction) and is experimentally supported by our data showing that CFM consolidation does not occur in the absence of sleep (Figure 2) – unless a network oscillation is artificially generated in CA1 (18). A simplified schematic of this idea is shown in Figure 6. While the present study is focused on matching parameters of computational models to data from the hippocampus during fear memory consolidation, we believe that the mechanisms outlined here may be universally true. For example, sleep, and sleep-associated network oscillations, are required for consolidation of experience-dependent sensory plasticity in the visual cortex (28, 32, 33). Moreover, similar frequency-dependent changes in neuronal firing rates are also observed across periods of sleep in the visual cortex (5). Based on these and other recent data linking network oscillations in sleep to many forms of memory consolidation, this suggests a unifying principle for sleep effects on cognitive function, and one that could reconcile discrepant findings on how sleep affects synaptic strength (15). It also paints a more complex but complete role of sleep in memory management that is often proposed (34). Here we show that sleep may on one hand facilitate disconnection of high frequency cells that initially participate in memory coding, but at the same time, mediates recruitment of new neuronal populations during consolidation of memory engram.

**Figure 6:**
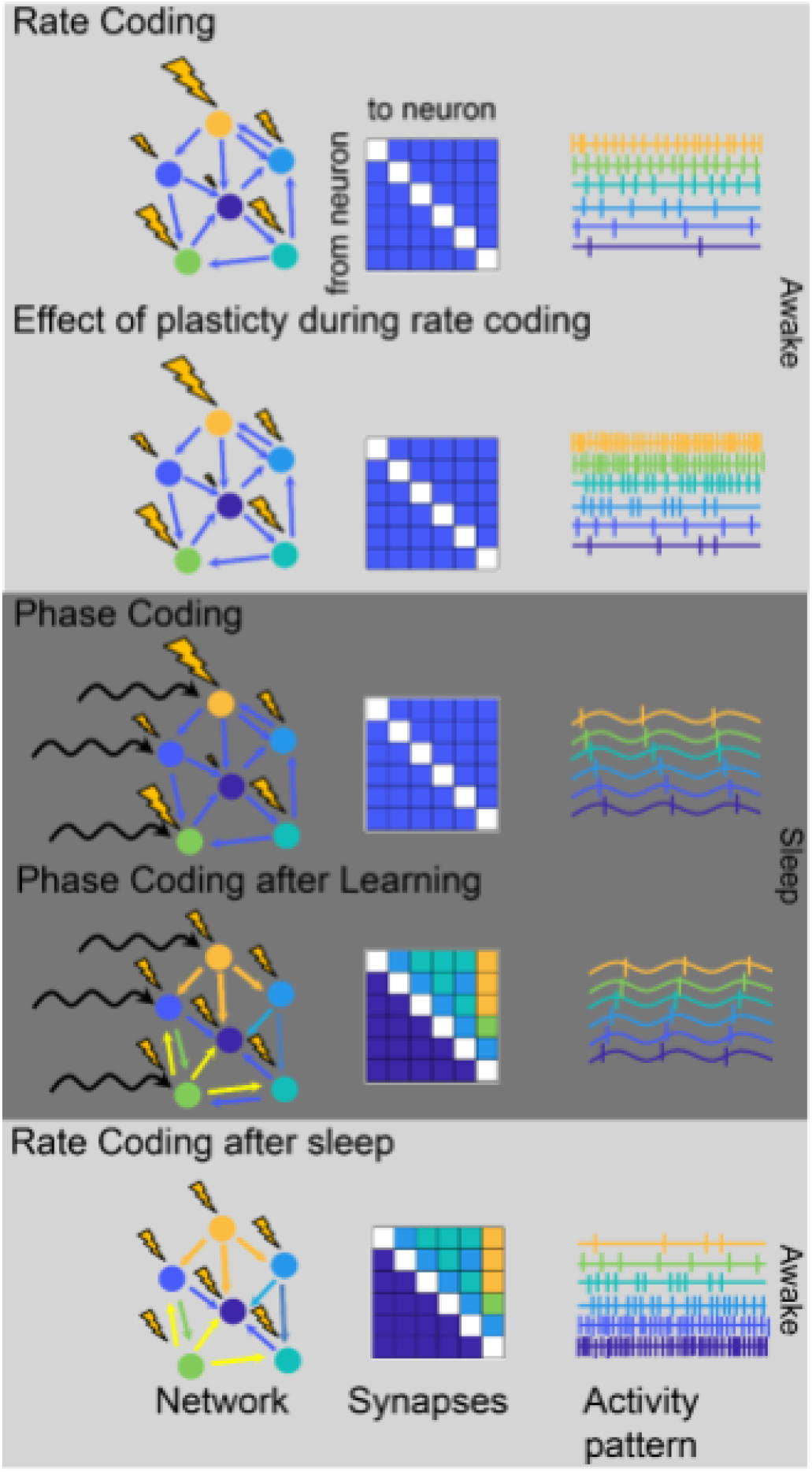
Phase coding and not rate coding explains memory consolidation and replay. Top (wake): Rate coding during wakefulness arranges spiking behavior in order of frequency, where neurons receiving a larger input fire faster. Learning during rate coding effectively speeds up the firing frequency and causes a uniform increase in all synaptic connectivity strengths. Middle (Sleep): Phase coding, on the other hand, occurs when neurons are firing at nearly the same frequency, with those neurons receiving greater input occurring first in a firing sequence. The effect of learning during rate coding is a preferential increase in synaptic connectivity strength, as well as de novo creation of new synapses, following the sequence. The firing phases eventually even out and then switch, so that the neurons firing first in the original sequence fire simultaneously and then last as learning progresses, respectively. Switching back to rate-coding after consolidation occurs during phase-coding effectively switches the order of the fastest firing neurons. Thereby, the process repeats itself and those neurons that are phase locked will see a net potentiation and therefore consolidation of a fear memory.

## Methods

### Mixed excitatory-inhibitory conductance-based neuronal networks

Conductance-based neuronal networks containing both excitatory and inhibitory neurons were modeled using a modified Hodgkin-Huxley formalism (21, 35). The time-dependent voltage *V*_*i*_ of a single neuron is given by

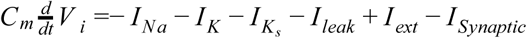

where *C*_*m*_ is the membrane capacitance, *I*_*ext*_ is the fixed external input used to elicit spiking, *I*_*leak*_ = 0.02(*V*_*i*_ + 60) is the leakage current, and 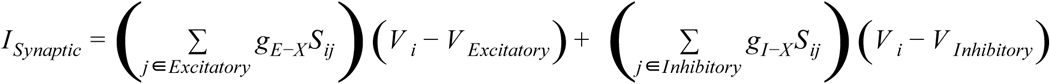 is the total summed synaptic input received by a neuron from its pre-synaptic partners and *g*_*I−X*_ and *g*_*E−X*_ represent the synaptic conductance for connections from inhibitory and excitatory neurons to their post synaptic targets *X* (values provided below). The synaptic reversal potentials are *V*_*Excitatory*_ = 0 *mV* and *V*_*Inhibitory*_ =− 75 *mV*. Here, 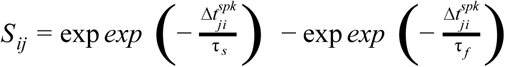 represents the shape of the synaptic current, given the difference in spike timing between the post-synaptic neuron *i* and the recently fired pre-synaptic neuron *j*, 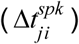, with τ_*f*_ = 5 *ms* and τ_*s*_ = 250 *ms* or τ_*s*_ = 30 *ms* for excitatory synaptic currents and inhibitory synaptic currents, respectively.

The ionic currents are *I*_*Na*_, *I*_*K*_, and *I*_*K*_*s*__, representing sodium (Na), potassium (K), and muscarinic slow potassium (K_s_), respectively. More specifically: 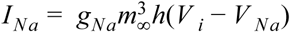, with 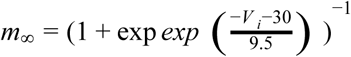 being the activation of the channel and where *h*, the inactivation, is given by the solution to 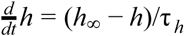, with 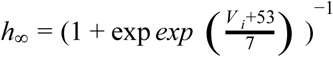 and 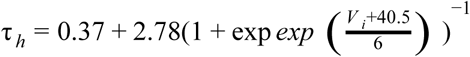; 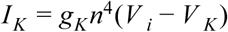 with 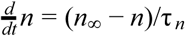 where 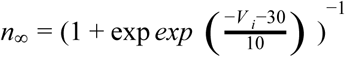 and 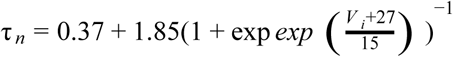 and 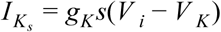 with 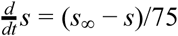 where 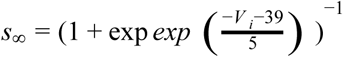. The reversal potentials are *V*_*Na*_ = 55 *mV* and *V*_*K*_ =− 90 *mV* and the maximal conductances are 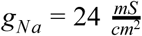, 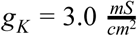.

The slow potassium conductance

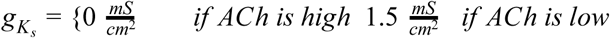

Controls the level of Acetylcholine (ACh), e.g. during wakefulness (high ACh) or NREM sleep (low ACh). The values thus control the excitability type, where low g_Ks_ (high ACh) yields Type 1 excitability and high gKs (low ACh) yields Type 2 excitability. Type 1 excitability is characterized by arbitrarily low firing frequencies, high frequency gain as a function of constant input, and a constant advance in the phase response curve whereas Type 2 has a threshold in firing frequency onset, a shallow frequency gain function,, and a biphasic phase response curve. See Stiefel, Gutkin, and Sejnowski (21).

Each simulation was completed using the RK4 integration method with a step size of h = 0.05 ms.

### Network properties

The network used in these studies consists of N=1000 neurons, with *N*_*e*_ = 800 excitatory neurons and *N*_*I*_ = 200 inhibitory neurons. Connections form a random network with different levels of connectivity dependent on the pairwise pre-and post-synaptic neuron identity: Inhibitory neurons project to 50% of the inhibitory neurons and 30% to the excitatory neurons whereas excitatory neurons project to just 6% of both the inhibitory and excitatory neurons, with self-connections being forbidden in all cases. The initial synaptic weights are *g*_*I−I*_ = 0.0013 *mS*/*cm*2, *g*_*I−E*_ = 0.0005 *mS*/*cm*2, 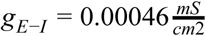, and *g*_*E−E*_ = 0.00003 *mS*/*cm*2 in Figures 1 and 2, but with *g*_*E−E*_ = 0.00001 *mS*/*cm*2 in Figures 3-5. For initial engram formation, a random subset of 250 excitatory neurons increase their *g*_*E−E*_ conductances, constituting the strength of the memory (referred to as synaptic heterogeneity). The effect of the increase in this synaptic strength is investigated in Figure 1 and subsequently kept at 10x in Figures 2-5.

### Learning through STDP

Neural correlates of memory are thought to emerge due to the strengthening and weakening of synaptic strengths in an activity-based manner following spike timing-dependent plasticity (STDP). Here, we use a symmetric learning rule that uniformly increases or decreases synaptic weights based on the time-ordering of pre-and post-synaptic pair firings, only in excitatory-to-excitatory connections. If a pre-synaptic neuron fires before its post-synaptic partner, the conductance increases by an amount 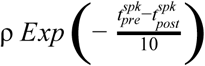. Similarly, a weakening of synaptic strength occurs by an amount 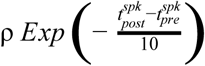 when a post-synaptic neuron fires before its pre-synaptic partner. In both cases, if the time difference between spike pairs is too great, the change in synaptic strength will approach zero. On the other hand, highly coincident spike pairs will have a maximal change given by the learning rate ρ = 10^−3^. It should be noted that while the synaptic weight is prohibited from becoming negative, there is no upper-bound set on the synaptic strength, though previous work has shown saturation of synaptic weights given sufficient time^14^.

### Hippocampal recordings, fear conditioning, and sleep deprivation

All procedures were approved by the University of Michigan Animal Care and Use Committee. Male C57BL/6J mice between 2 and 6 months were implanted using methods described previously (33, 36). The implants (described in more detail in Ognjanovski N et al. [17]) consisted of custom built driveable headstages with two bundles of stereotrodes implanted within right-hemisphere CA1. 3 EMG electrodes were placed in nuchal muscle to monitor activity. The signals from the stereotrodes were split into local field potential (0.5-200 Hz) and spike data (200 Hz-8 kHz). Single neuron data was discriminated as described in Ognjanovski N et al. (17), and only neurons stably recorded across each 24 hour period were used in the subsequent analyses.

After surgical recovery, the animals were recorded for a 24 hour baseline. The next day, following procedures described here, the animals underwent either a shock or no shock (sham experience). Post-shock, animals were either allowed ad lib sleep or 6 hours of sleep deprivation via gentle handling. 24 hours later, freezing behavior to conditioning context was assessed to evaluate the formation of a contextual fear memory.

### Pharmacogenetic inhibition of interneurons

2-3 month old male Pvalb-IRES-CRE mice were bilaterly injected with either the inhibitory receptor hM4Di (rAAV2/Ef1A-DIo-hM4Di-mCherry; UNC Vector Core: Lot #AV4,708) or a control mCherry reporter (raav2/Ef1A-DIo-mCherry; UNC Vector Core: Lot #AV4375FA) (methods further elaborated in Ognjanovski N et al. [18]). Using the same implant procedures described above, the animals were implanted with stereotrode bundles.

After allowing 4 weeks for viral expression, the animals underwent contextual fear conditioning (as described above). Post-shock, mice were either given 0.04 mL i.p. injecting of either 0.3 mg/kg clozapine-N-oxide dissolved in DMSO (to activate the DREADD) or DMSO alone (as a control). Further detail can be found in Ognjanovski N et al. (20), where this data set was originally published.

### Optogenetic inhibition and stimulation

Pvalb-IRES-CRE mice were crossed with B6.Cg-Gt(ROSA)^26Sortm40.1(CAG-aop3/EGFP)Hze^/J, B6;129S-Gt[ROSA]26Sor^tm32(CAG-COP4*H134R/ EYFP)Hze^/J, or B6.Cg-Gt(ROSA)26Sor^tm6(CAG-ZsGreen1)Hze^/J transgenic mice. This gave 3 groups of mice – one expressing Arch in parvalbumin interneurons (PV::Arch), one expressing ChR2 in parvalbumin interneurons (PV::ChR2), and one expressing eGFP in parvalbumin interneurons (PV::GFP). Using similar implant procedures wo above, the animals were bilaterally implanted with stereotrode bundles in CA1 in addition to optical fibers to deliver laser light.

Following postoperative recovery, the animals underwent contextual fear conditioning as described above. PV::Arch mice received state-specific inhibition of parvalbumin interneurons using 3 mW 532 nm laser light. The data used in this paper focuses on the NREM specific inhibition. PV::GFP mice received the same state-specific laser light as a control. PV:ChR2 mice were sleep deprived post-conditioning while simultaneous receiving 20 ms square-wave pulses of 473 nm light at 7 Hz (estimated power 3-10 mW). PV::GFP controls underwent the same sleep deprivation and light pulses.

### Analysis of functional network structures through AMD and FuNS

Average Minimal Distance (AMD) (37) was applied to network spiking data to determine functional connectivity. AMD calculates the mean value of the smallest temporal difference between all spikes in one neuron and all spikes in another neuron. Analytical calculations of the expected mean and standard deviation of minimal distance is then used to rapidly determine the significance of pairwise minimal distance (24). Specifically, the first and second raw moments of minimal distance for each node are calculated: 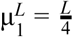 and 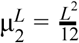, where L is the temporal length of the interspike interval and we have assumed that (looking both forward and backward in time) the maximum temporal distance between spikes is 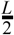. Over the entire recording interval T, the probability of observing an inter-spike interval of length L is simply 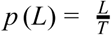. Then, the first and second moments of minimal distance considering the full recording interval are given as 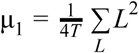 and 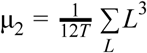. Finally, the calculated statistical moments give rise to the expected mean and standard deviation, μ = μ_1_ and 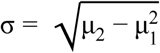, which are used to determine the Z-score significance of pairwise connectivity: 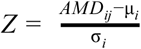. Values of *Z*_*ij*_≥2 represent significant functional connections between node pairs.

Functional Network Stability (FuNS) tracks global changes in network functional connectivity by quantifying similarities in AMD matrices over a recording interval (24). The procedure is as follows: first, a recording interval is split into n partitions of equal temporal length. Each partition is subjected to AMD functional connectivity analysis, resulting in n functional connectivity matrices Z. Similarity between time-adjacent functional networks is determined using the normalized dot product after matrix vectorization. FuNS is then determined by taking the mean of these cosine similarities over the recording interval: 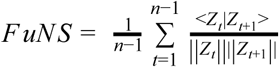. Thus, not only does FuNS yield insight into how functional connectivity changes over time, it can also shed light on how behavior affects the underlying functional network, e.g. by quantifying the difference between FuNS calculated before and after a learning task.

### Spectral Analysis, spike-field coherence, and phase relationships

Histograms of neuronal spiking per unit time were used to calculate the network characteristic frequency, spike-field coherence, and phase relationships of individual neurons to the network signal. First, spike timings were converted into binary spike vectors and then summed to give a network spike vector. Then, the spike vector was convolved with a Gaussian distribution with zero mean and a standard deviation of ~2 ms, giving a continuous network signal. The spectral power was measured by taking the Fourier transform of the excitatory, non-engram signal (i.e. no neurons with artificially strengthened connections were used). Then, the change in spectral power (e.g., in Figure 1) was calculated by integrating the frequency-domain signals and taking the relevant percent difference.

Conversely, the spike-field coherence was measured by comparing the non-inhibitory, non-engram network signal to the individual excitatory signals making up the general signal (i.e. convolving an individual neuron’s spike vector) by taking the cross correlation at zero lag. The values reported were normalized using the autocorrelation, giving values between 0 (no correlation) and 1 (perfect correlation).

Finally, the phase relationships reported were calculated using a continuous inhibitory signal from the inhibitory neuron population spike vector. Peaks in the inhibitory signal acted as the start and end of a given phase and non-engram excitatory spike times were used to calculate the phase difference between an individual neuron and the signal. The phase difference was normalized to give values between 0 and 1.

